# DeepRaccess: High-speed RNA accessibility prediction using deep learning

**DOI:** 10.1101/2023.05.25.542237

**Authors:** Kaisei Hara, Natsuki Iwano, Tsukasa Fukunaga, Michiaki Hamada

**Affiliations:** Department of Electrical Engineering and Bioscience, Graduate School of Advanced Science and Engineering, Waseda University, Tokyo, 1698555, Japan; Computational Bio Big-Data Open Innovation Laboratory, AIST-Waseda University, Tokyo, 1698555, Japan; Waseda Institute for Advanced Study, Waseda University, Tokyo, 1690051, Japan; Graduate School of Medicine, Nippon Medical School, Tokyo 1138602, Japan

## Abstract

RNA accessibility is a useful RNA secondary structural feature for predicting RNA-RNA interactions and translation efficiency in prokaryotes. However, conventional accessibility calculation tools, such as Raccess, are computationally expensive and require considerable computational time to perform transcriptome-scale analyses. In this study, we developed DeepRaccess, which predicts RNA accessibility based on deep learning methods. DeepRaccess was trained to take artificial RNA sequences as input and to predict the accessibility of these sequences as calculated by Raccess. Simulation and empirical dataset analyses showed that the accessibility predicted by DeepRaccess was highly correlated with the accessibility calculated by Raccess. In addition, we confirmed that DeepRaccess can predict protein abundance in *E.coli* with moderate accuracy from the sequences around the start codon. We also demonstrated that DeepRaccess achieved tens to hundreds of times software speed-up in a GPU environment. The source codes and the trained models of DeepRaccess are freely available at https://github.com/hmdlab/DeepRaccess.

## Introduction

RNA molecules play crucial roles in the regulation of diverse cellular processes, and their regulatory functions are closely linked to their structures ([1]). For example, tRNAs have to form cloverleaf secondary structures and L-shaped tertiary structures in order to function properly during translation. As another example, in prokaryotic translation, the RNA region upstream of the start codon has a function to regulate protein abundance, and the level of abundance decreases when the region takes a stem structure ([2]). Accordingly, many experimental and computational studies have been carried out to analyse RNA structures in order to elucidate the relationships between the structures and functions ([3, 4]). In particular, computational analyses of RNA secondary structures are frequently performed because of their low cost, moderate accuracy, and high speed ([5, 6, 7, 8, 9]).

RNA accessibility is one of the secondary structural features and is defined as the energy required for an RNA region not to form a stem structure. The accessibility is used to predict RNA-RNA interactions ([10, 11, 12]) and translation efficiency in prokaryotes ([13]) because these molecular processes are more likely to occur when the RNA region of interest is single-stranded. Therefore, several software programs have been developed to calculate the RNA accessibility ([14, 15, 16, 17, 18]). Some of these programs used a local folding approach, which reduces computational time by ignoring long-distance base pairs ([14, 17, 18]). However, current methods are still too computationally expensive for transcriptome-scale analysis, and thus the development of faster methods for calculating accessibility is an essential research topic. In general, one of the powerful approaches to speed up the calculation is parallel computing, and several parallel algorithms have now been proposed for RNA secondary structure analysis ([19, 20]). However, because most algorithms for RNA secondary structure analysis are based on dynamic programming, which is difficult to parallelize, the parallel algorithms have not been fully explored, especially in parallel computations using GPUs ([21]).

In recent years, machine learning-based simulation acceleration has attracted attention in computer simulation ([22, 23]). This method uses running results of a slow but accurate simulator as training data, and constructs a predictive model that reproduces the simulation results. Since the run of the predictive model is generally much faster than that of the simulator, the accurate predictive model can be seen as a fast alternative to the simulator. In particular, deep learning-based methods have the advantage of using GPUs efficiently based on the deep learning libraries without the need to build specialised algorithms. Machine learning-based acceleration is beginning to be used in bioinformatics, such as phylogenetic tree construction ([24]) and sequence alignment score calculation ([25, 26, 27, 28]). However, there is no research on the application to RNA secondary structure analysis.

In this study, we developed DeepRaccess, a fast accessibility prediction tool based on deep learning-based software acceleration. We confirmed that DeepRaccess can moderately reproduce the results of Raccess, an existing RNA accessibility calculation method ([17]), with high accuracy on both simulation and empirical datasets. We also demonstrated that the accessibility calculated by DeepRaccess is moderately correlated with protein abundance in *E. coli*. Finally, we verified that DeepRaccess was significantly faster than Raccess on various datasets in a GPU environment.

## Methods

### Overview of the DeepRaccess software

DeepRaccess is a machine learning predictor whose input is an RNA sequence and whose output is the accessibility in subregions of the sequence. Fig. 1 shows an overview of the DeepRaccess approach. The subregion length *l*_*a*_ is fixed in the training step, and the accessibility of all subregions with the length *l*_*a*_ are the output. When users require the accessibility with a different length *l*_*a*_, they have to redo the training of the prediction model. In this study, we used 35 as the default value for *l*_*a*_. Note that this value has been used in a previous study to predict prokaryotic translation efficiency ([13]).

**Figure 1:**
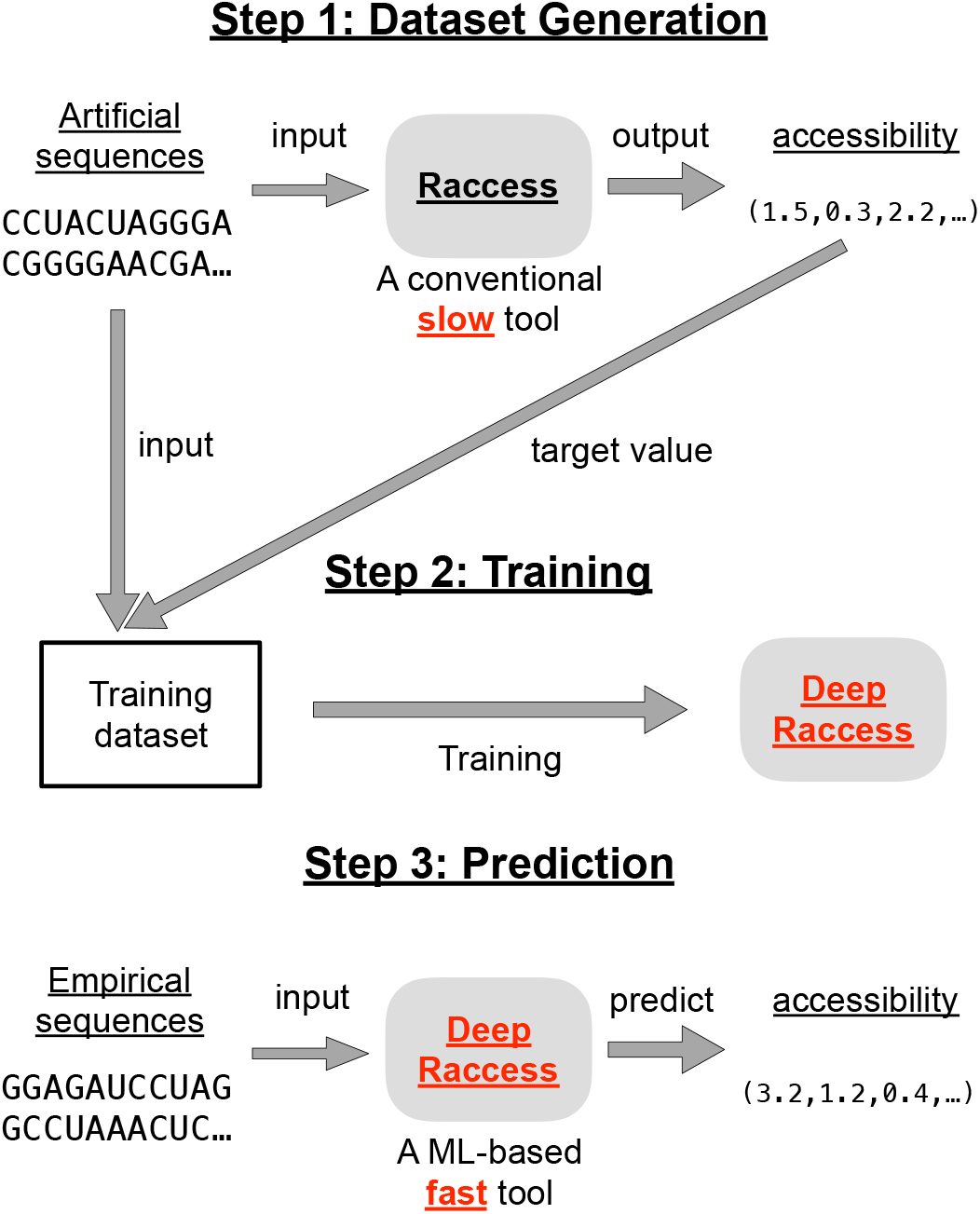
The schematic illustration of the DeepRaccess approach. DeepRaccess first trains a model to predict accessibility from RNA sequences by utilizing a training dataset composed of artificial RNA sequences and their corresponding accessibility calculated by Raccess. DeepRaccess then rapidly predicts the accessibility of empirical RNA sequences based on the trained predictive model.

The training datasets consisted of RNA sequences as the input and the accessibility as the target values. The RNA sequences were artificially generated (the details are described in Section 2.3), and the accessibility was calculated from the input RNA sequences using Raccess ([17]). Raccess adopts a local folding approach that speeds up the computation by ignoring base pairs spanning more than *W* bases, and can compute the accessibility of all subregions based on a secondary structure score model for a fixed *l*_*a*_ length. The computation is based on dynamic programming, and the time complexity is *O*(*NW*^2^) where *N* is the sequence length. In this study, we used the CONTRAfold model as the score model because of its high accuracy ([29]), and used 100 as the default value of *W*. Note that Raccess is the only software that can compute the accessibility for long sequences in a reasonable time under the numerically stable computation. The source code for Raccess is not available from the link given in the original Raccess paper, but can be downloaded from the following link: https://github.com/gterai/raccess.

Because we set the maximum sequence length in the training dataset to 440, which is the value used in RNABERT ([30]), we could not predict the accessibility of sequences longer than 440 bases by merely applying DeepRaccess without any modifications. Therefore, for such a long sequence, we used DeepRaccess by implementing the following procedures. First, DeepRaccess divides the sequence into subsequences of 440 bases by shifting the window by 330 bases. This means that neighboring subsequences overlap by 110 bases. DeepRaccess then predicted the accessibility of these subsequences and integrated them with the accessibility of the full-length RNA. In the overlapped region, DeepRaccess ignored the accessibility of the 55-base region from the end of each subsequence.

### Neural network architecture

We used deep neural networks as the predictor. We implemented four representative network architectures using PyTorch and compared their prediction accuracy: (1) Fully Convolutional Network (FCN) ([31]), (2) U-Net ([32]), (3) Bidirectional Encoder Representations from Transformers (BERT) ([33]) and (4) RNABERT ([30]). Tables S1–4 show the details of each network architecture, respectively. We set the epoch and batch sizes to ten and 256, respectively. We also used AdamW as the optimizer and the values used by RNABERT as the hyperparameters of the optimizer.

The input RNA sequences are embedded into numerical vectors and fed into the neural networks. The FCN and U-Net models used token embedding. This embedding first randomly generates six 120-dimensional numerical vectors corresponding to each of the six states: four RNA bases (A, C, G, U), one undetermined nucleotide (N) and padding. The resulting vectors are then assigned to each state in the input sequences. The BERT and RNABERT models used positional embedding in addition to token embedding. In this embedding, 120-dimensional numerical vectors corresponding to each position in the sequences are randomly generated. Finally, the values of the two embedding results are summed for each base.

We briefly review each network architecture. FCN is a type of CNN architecture that is widely used for image segmentation, and the characteristic is that it does not use fully connected layers and is composed only of convolutional layers. We used a network of 40 convolutional layers with constant channel and unit sizes as the FCN model. U-Net is a variant of FCN, and consists of three parts, bottom-up path, bottleneck, and top-down path. The data is downsampled in the bottom-up path, the computation is performed in convolutional layers with the smallest unit sizes in the bottleneck, and the data is upsampled in the topdown path. The essential feature of U-Net is that the layers on the bottom-up and top-down paths have skip connections. For the U-Net model, we used a network consisting of 3, 35, and 3 layers on the bottom-up path, bottleneck, and top-down path, respectively. BERT was originally developed for natural language processing and is a model in which transformer layers are stacked several times. In this study, we stacked six transformer layers. Transformer can incorporate positional information of elements into the model by using the attention mechanism. RNABERT is a BERT model pre-trained on 76,237 human small ncRNAs in the RNAcentral database ([34]), and both RNA sequence and structural information were embedded in the learned representation of RNABERT.

### Training Datasets

All sequence data used for training were artificially generated. We generated the sequences using two methods: 1) uniform base sampling to generate RNAs that lack strong stem structures and 2) sampling to promote the formation of stem structures to generate RNAs with strong stem structures similar to small ncRNAs. In this paper, we refer to these methods as the uniform and the structured RNA sampling methods, respectively.

In the uniform sampling method, we first determined the sequence length *N* by sampling from the uniform distribution *uni f* (100, 440). The bases in the sequences were sampled from the categorical distribution *Cat*(*x*|*π*) for the category (A, C, G, U, N), and *π* was sampled from the Dirichlet distribution *Dir*(*π* |*α* = [1, 1, 1, 1, 0.1]). *π* was sampled once per sequence.

In the structured RNA sampling method, after generating a sequence based on the uniform sampling method, we determined the stem length *l* by sampling from *uni f* (8, 48). We next selected the length *d* of the region flanked between two stem regions from *uni f* (3, *N* − 2*l*) and the start position of the first stem region from *uni f* (0, *N* − 2*l* − *d*). We then substituted the bases in the second stem regions so that the bases were complementary to the bases of the first region. Here, when the base was G or U, whether it formed a Watson-Crick base pair or a wobble base pair was determined by the Bernoulli distribution *Bern*(*x*|*μ*), and *μ* was sampled from the Beta distribution *Beta*(*μ*|*α* = 4, *β* = 1). After that, we substituted the bases in the stem region to create internal loops, and whether a base was substituted or not was determined by *Bern*(*x*|*μ*). Here, *μ* was sampled from *Beta*(*μ*|*α* = 1, *β* = 15), and the base after the substitution was sampled from *Cat*(*x*|*π*) using the uniform sampling method. We also substituted the next base after the substituted base according to *Bern*(*x*|*μ*), and *μ* was sampled from *Beta*(*μ*|*α* = 2, *β* = 1).

Using these two methods, we created two training datasets that were a uniform RNA dataset and a structured RNA dataset. In the former, all sequences were generated by the uniform sampling method, while the latter contained half of each of the sequences generated by the two sampling methods. We performed the training on each of the two training datasets and created two predictive models for each architecture. We used 10 million as the default number of sequences per the training.

### Test datasets and evaluation measure

We evaluated the prediction accuracy of DeepRaccess using simulation test datasets and three empirical datasets: Rfam, Gencode, and *E.coli* synthetic mRNA datasets. As the test simulation dataset, we used a dataset generated in the same way as the training data used for the trained model. We used 100 thousand as the number of sequences per the test dataset. The Rfam dataset consisted of 3,105,149 sequences from the Rfam 14.9 database ([35]), and most of which are highly structured. The Gencode dataset contains 142,379 transcripts in Gencode version M29 ([36]). We have removed sequences of less than 35 bases from this analysis. The *E.coli* synthetic mRNA datasets consisted of 244,000 sequences with 120 bases around the start codon of synthetic mRNAs ([37, 13]). We applied Raccess and DeepRaccess to these datasets, and then compared the accessibility calculated by the methods. For the evaluation measure, we used Spearman’s rank correlation coefficient (*ρ*) and normalised mean square error (NMSE), which is the MSE divided by the target value.

To validate the usefulness of DeepRacess, we also evaluated its predictive performance for prokaryotic translation efficiency. We used the *E. coli* synthetic mRNA dataset for this analysis, and calculated Spearman’s *ρ* between the accessibility based on the DeepRaccess and protein abundance. For the comparison, we used the accessibility calculated by Raccess, minimum free energy (MFE) calculated by CONTRAfold ([29]), and scores of RBSDesigner ([38]) and RBSCalculator ([39]).

We investigated the computational speed of DeepRaccess and compared it to Raccess. Raccess and DeepRaccess were run in a CPU-only environment (CPU: Intel(R) Xeon(R) Gold 6148 2.1 GHz, memory:8GB). In addition, DeepRaccess was also run in an environment where both CPU and GPU were available (CPU: Intel(R) Xeon(R) CPU E5-2698 v4 2.2 GHz, GPU: Tesla V100 DGXS 32GB×4, memory: 257GiB).

## Results

### Accuracy evaluation on simulation datasets

We first evaluated the prediction accuracy of DeepRaccess using simulation test datasets and compared the performances of different neural network architectures. Table 1, Fig. 2, and Fig. S1 show the prediction performances of DeepRaccess. We found that the NMSEs were less than 0.25 and the Spearman’s *ρ*s were greater than 0.97 in all cases, suggesting that deep learning is effective in predicting RNA accessibility. In addition, the scores based on the structured RNA dataset were worse than those based on the uniform RNA dataset in each architecture. The reason for the difficulty in prediction may be that the structured RNA dataset has a large variance in the RNA accessibility. We also verified that the FCN was the best-performing architecture in both datasets and therefore used the FCN in the following analyses.

**Table 1:**
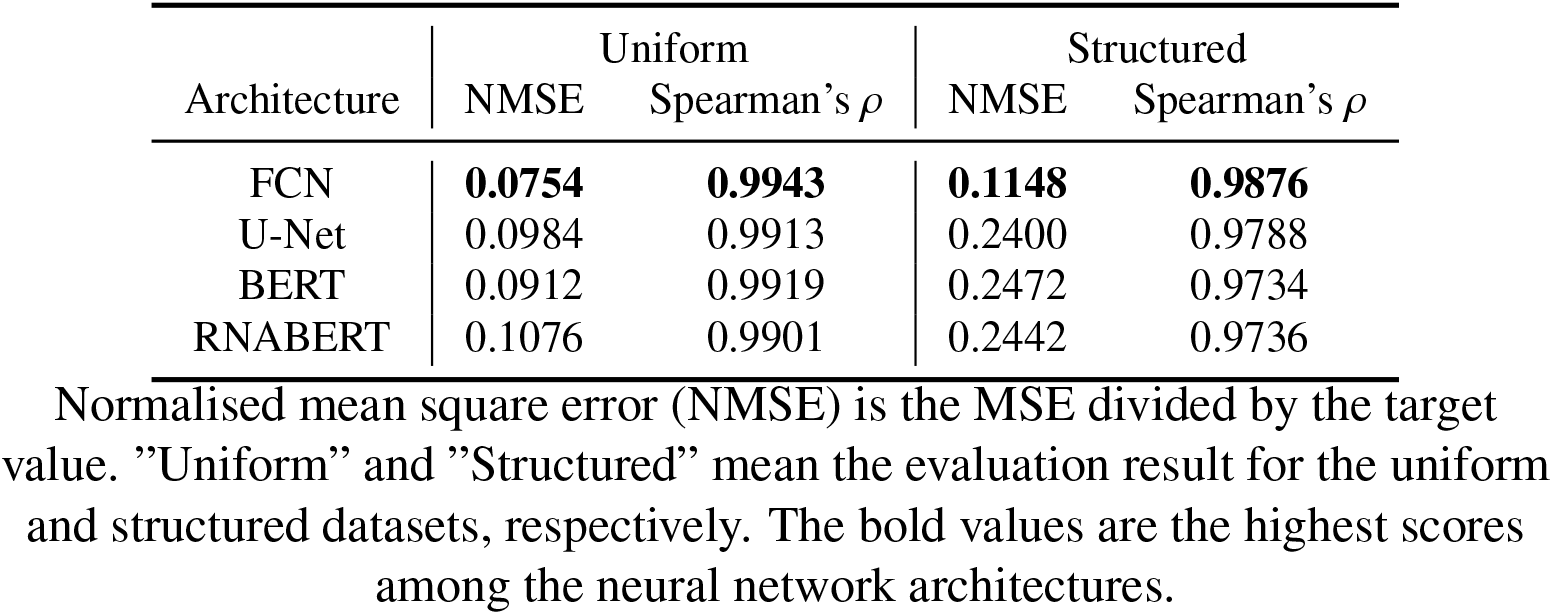
Comparison of the prediction accuracy among the neural network architectures

| Architecture | Uniform |  | Structured |  |
| --- | --- | --- | --- | --- |
| | NMSE | Spearman's $\rho$ | NMSE | Spearman's $\rho$ |
| FCN | **0.0754** | **0.9943** | **0.1148** | **0.9876** |
| U-Net | 0.0984 | 0.9913 | 0.2400 | 0.9788 |
| BERT | 0.0912 | 0.9919 | 0.2472 | 0.9734 |
| RNABERT | 0.1076 | 0.9901 | 0.2442 | 0.9736 |

Normalised mean square error (NMSE) is the MSE divided by the target value. "Uniform" and "Structured" mean the evaluation result for the uniform and structured datasets, respectively. The bold values are the highest scores among the neural network architectures.

**Figure 2:**
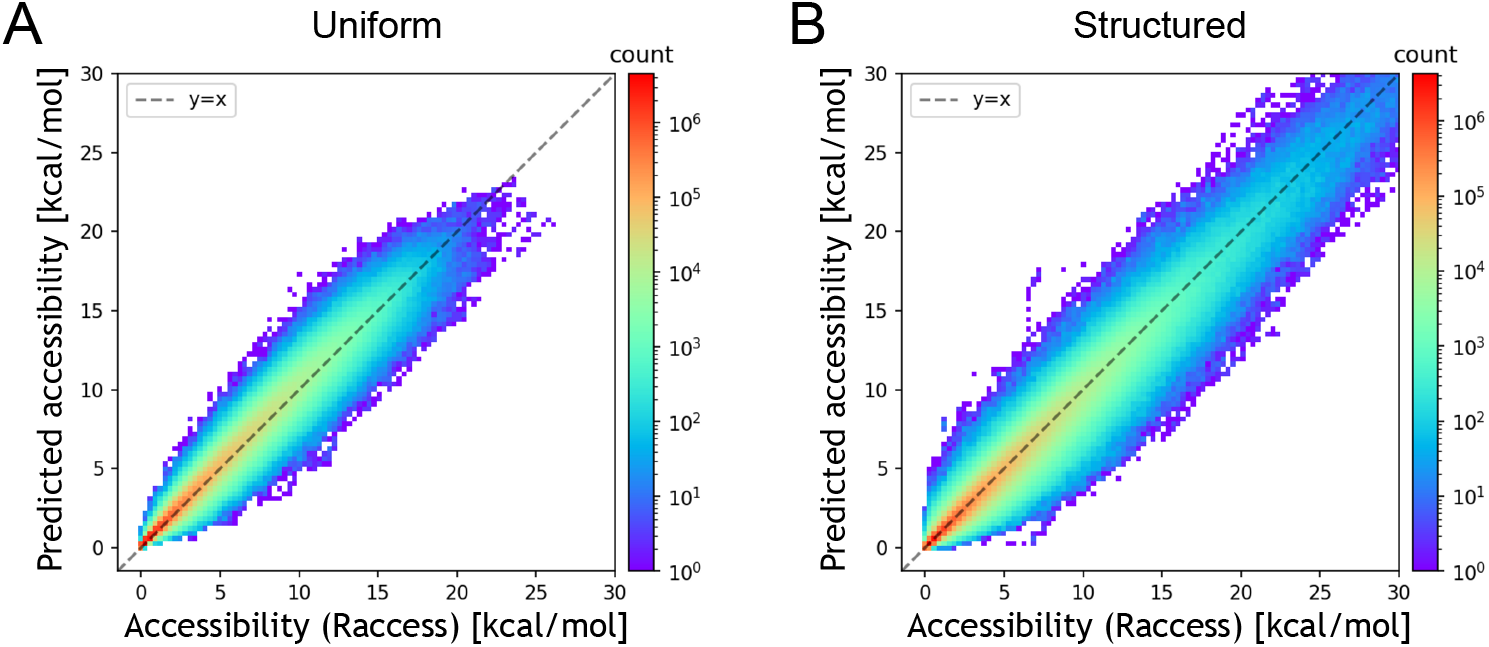
Prediction accuracy of the FCN architecture for the simulation datasets ((A) the uniform RNA dataset and (B) the structured RNA dataset). The x and y axes represent the accessibility calculated by Raccess and predicted accessibility, respectively. The color bar representing the counts is displayed using a log scale.

We next investigated the effect of the training data size on the prediction performances using the structured RNA dataset (Table S5 and Fig. S2). We confirmed that the prediction accuracy improved as the data size increased, and the accuracy had not yet converged even when the data size was increased to 10 million. Therefore, we should achieve higher prediction accuracy as more datasets are prepared and more time is spent on training. We also evaluated the effect of the parameters *l*_*a*_ and *W* on the performances (Table S6 and Fig. S3). We found that DeepRaccess had higher performance when *l*_*a*_ was large. In addition, small *W* resulted in accurate prediction. This may be because the RNA accessibility has small variances and ranges when *W* is small.

### Accuracy evaluation on the empirical datasets

We then assessed whether DeepRaccess could predict the accessibility of empirical RNA sequences using three datasets (Table 2, Fig. 3, and Fig. S4). We verified that the best predictor for each dataset has an NMSE of less than 0.23, and the Spearman’s *ρ* was greater than 0.88 for all datasets. While the predictive model trained on the uniform RNA dataset outperformed that trained on the structured RNA dataset for the Gencode and *E.coli* synthetic mRNA datasets, the opposite trend was found for the Rfam dataset. In addition, the prediction results on the Rfam dataset had the highest NMSE of the three datasets. These results are probably due to the fact that the Rfam dataset contains many structured RNAs. Furthermore, we found some data on the Rfam and Gencode datasets where the predicted accessibility was very low when the accessibility calculated by Raccess was large. This result means that these regions were predicted to form almost no stems, despite actually having strong stem structures. In conclusion, DeepRaccess was also able to predict RNA accessibility with high accuracy for empirical RNA sequences, but its accuracy was insufficient for highly structured RNAs.

**Table 2:** Prediction performances for the empirical datasets

| Dataset | Uniform |  | Structured |  |
| --- | --- | --- | --- | --- |
| | NMSE | Spearman's $\rho$ | NMSE | Spearman's $\rho$ |
| Gencode | 0.1352 | 0.9215 | 0.1422 | 0.9159 |
| Rfam | 0.5493 | 0.8721 | 0.2244 | 0.9044 |
| *E.coli* | 0.1186 | 0.8821 | 0.1520 | 0.8548 |

**Figure 3:**
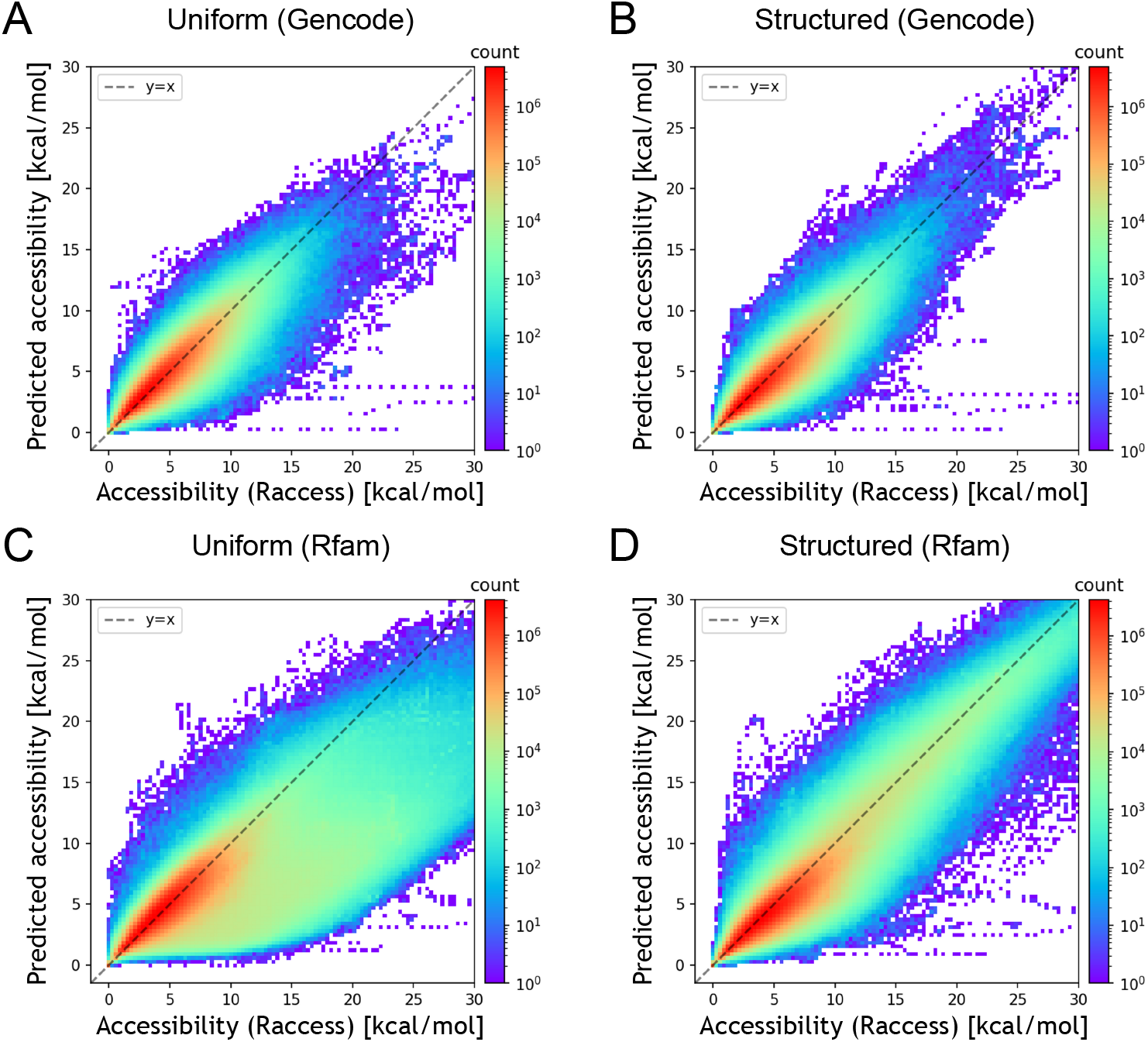
Prediction accuracy for the empirical datasets. (A) Prediction performances for the Gencode dataset of the predictive model trained on the uniform RNA dataset and (B) the structured RNA dataset. (C) Prediction performances for the Rfam dataset of the predictive model trained on the uniform RNA dataset and (D) the structured RNA dataset. The x and y axes represent the accessibility calculated by Raccess and predicted accessibility, respectively. The color bar representing the counts is displayed using a log scale.

We also evaluated the correlation between the protein abundance in *E.coli* and the accessibility calculated by DeepRaccess (Table 3 and Fig. 4). We used the predictive model trained on the uniform RNA dataset because of the high accuracy. We found that the Spearman’s *ρ* was 0.585, indicating that DeepRaccess could predict the protein abundance with moderate accuracy. The prediction accuracy of DeepRaccess was lower than that based on the accessibility calculated by Raccess, but comparable to that of the MFE score and higher than those of RBSDesigner and RBSCalculator.

**Table 3:**
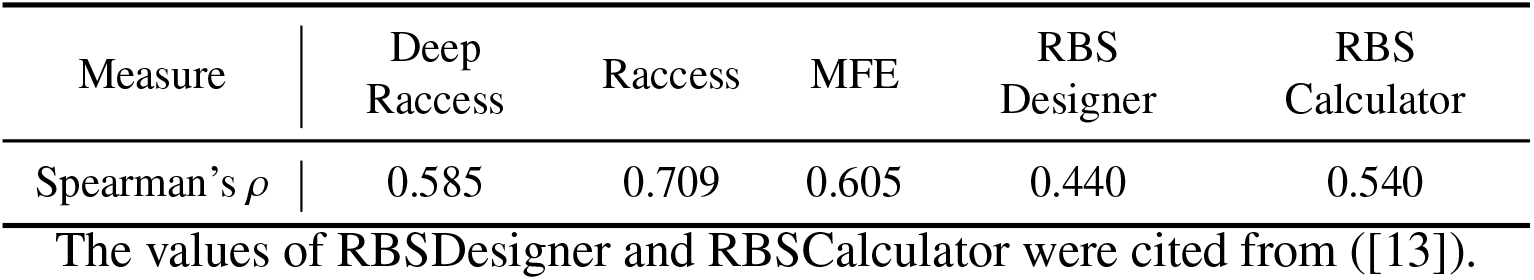
Prediction performances for the protein abundance in the *E.coli* synthetic mRNA dataset

| Measure | Deep<br>Raccess | Raccess | MFE | RBS<br>Designer | RBS<br>Calculator |
| --- | --- | --- | --- | --- | --- |
| Spearman's $\rho$ | 0.585 | 0.709 | 0.605 | 0.440 | 0.540 |

The values of RBSDesigner and RBSCalculator were cited from ([13]).

**Figure 4:**
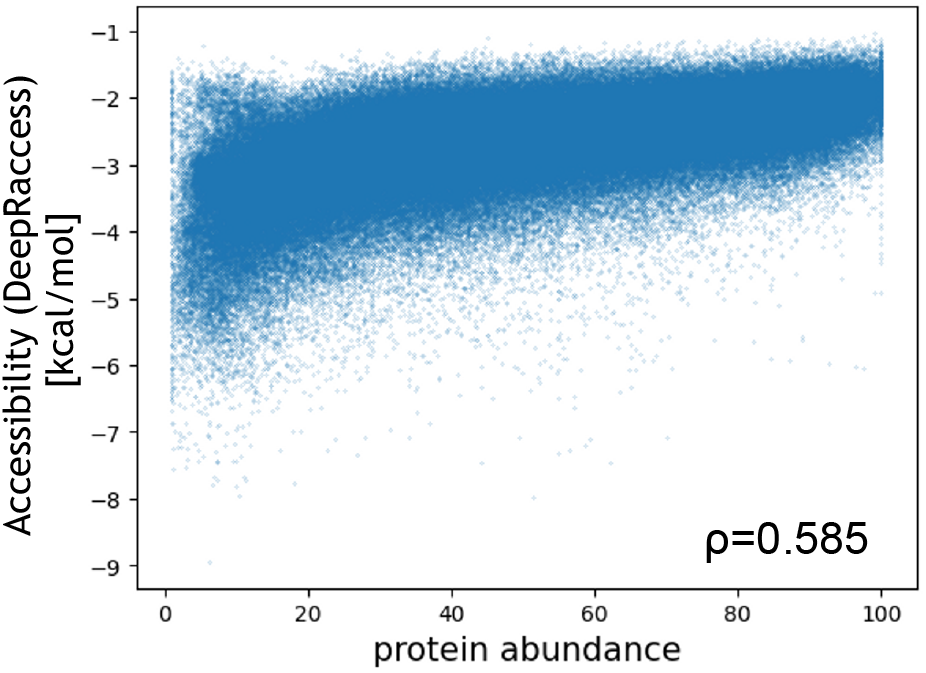
Correlation between the protein abundance and the accessibility calculated by DeepRaccess. The protein abundance was measured by fluorescence-activated cell sorting and was normalized so that the minimum value is 1 and the maximum value is 100 ([37]). The x- and y-axis represent the protein abundance and the accessibility, respectively.

### Evaluation of the computational speed

Finally, we evaluated the runtime of DeepRaccess. First, we looked at the time taken to train the predictive model and found it to be 2 days and 20 hours in the GPU environment. This is not short but the prediction model only needs to be trained once, and thus the long time is not a practical bottleneck. Note that this training step is not necessary when users are using trained models of DeepRaccess. We next assessed the time taken to predict the RNA accessibility (Table 4). In the CPU-only environment, DeepRaccess was not necessarily faster than Raccess, and the superiority depended on the datasets. On the other hand, DeepRaccess was tens to hundreds of times faster in the GPU environment than Raccess in the CPU environment. Although it should be noted that the environment in which the computations were performed was different, we have shown that DeepRaccess was extremely fast compared to Raccess.

**Table 4:** The run time evaluation on simulation and empirical datasets

| Program | Simulation | Gencode | Rfam | *E.coli* |
| --- | --- | --- | --- | --- |
| Raccess | 11h52m<br>(628.3) | 5d22h<br>(82.0) | 6d01h<br>(183.5) | 8h53m<br>(201.1) |
| DeepRaccess:<br><CPU> | 3h02m<br>(160.3) | 9d03h<br>(126.5) | 4d23h<br>(149.9) | 8h24m<br>(190.1) |
| DeepRaccess:<br><GPU> | 1m08s<br>(1.0) | 1h44m<br>(1.0) | 47m33s<br>(1.0) | 2m39s<br>(1.0) |

The rows indicate the run times and the run time ratio of each program to DeepRaccess <GPU>.

## Discussion

In the current study, we proposed DeepRaccess, a rapid RNA accessibility prediction method based on the deep learning. We evaluated the prediction accuracy and the computational speed of DeepRaccess using the simulation and three empirical datasets, and validated that DeepRaccess had a high level prediction accuracy while exhibiting significantly faster performance on the GPU environment. In addition, we demonstrated that the accessibility of regions around start codons of *E.coli* mRNA calculated by DeepRaccess can predict the protein abundance.

Moreover, there is scope for improvement in training data generation methods. In this study, we employed two sampling methods: uniform and structured RNA sampling. However, utilizing enhanced sequence data generation methods that yield data more akin to empirical RNA sequences should lead to improved prediction accuracy. Deep generative models, such as generative adversarial networks and diffusion models, may hold promise as potential methods.

Although DeepRaccess had high prediction accuracy, further improvement in prediction performance is an essential issue. The simplest approach is to increase the number of training data. In this study, we used 10 million RNA sequences as our training data, but we expect to improve the accuracy by using several billion RNA sequences. While increasing data size is difficult in machine learning for bioinformatics in general, our method allows unlimited data growth by generating data through simulation. In addition, there is scope for improvement in training data generation methods. In this study, we employed two sampling methods: the uniform and structured RNA sampling methods. However, utilizing enhanced sequence data generation methods that produce data more akin to empirical RNA sequences should improve the prediction accuracy. Deep generative models, such as generative adversarial networks ([40]), should hold promise as potential methods. Furthermore, the development of neural network architectures is also a promising approach for improving accuracy. For example, Corso *et al*. proposed that embedding in the hyperbolic space improves the accuracy of predicting the edit distance between sequences ([26]), and thus applying the non-Euclidean space may also be useful in predicting RNA accessibility ([41]).

The computational speedup provided by the deep learning method can be applied to the other secondary structural features such as base pairing probabilities (([42])), structural profiles ([43]), and structural entropy ([44]). Each feature has been used to improve the accuracy of RNA secondary structure prediction ([45]), to predict RNA-protein binding ([46]), and to evaluate the effect of base mutations on the structure ([47]). In particular, the algorithm used to compute these structural features taking into account pseudoknots is extremely slow ([48]), and thus speeding up the method through deep learning should be an important topic of future research.

## Acknowledgments

Computations were performed on the NIG supercomputer at ROIS National Institute of Genetics.

