## Supplementary Materials for "DeepRaccess: High-speed RNA accessibility prediction using deep learning"

### Supplementary Figures

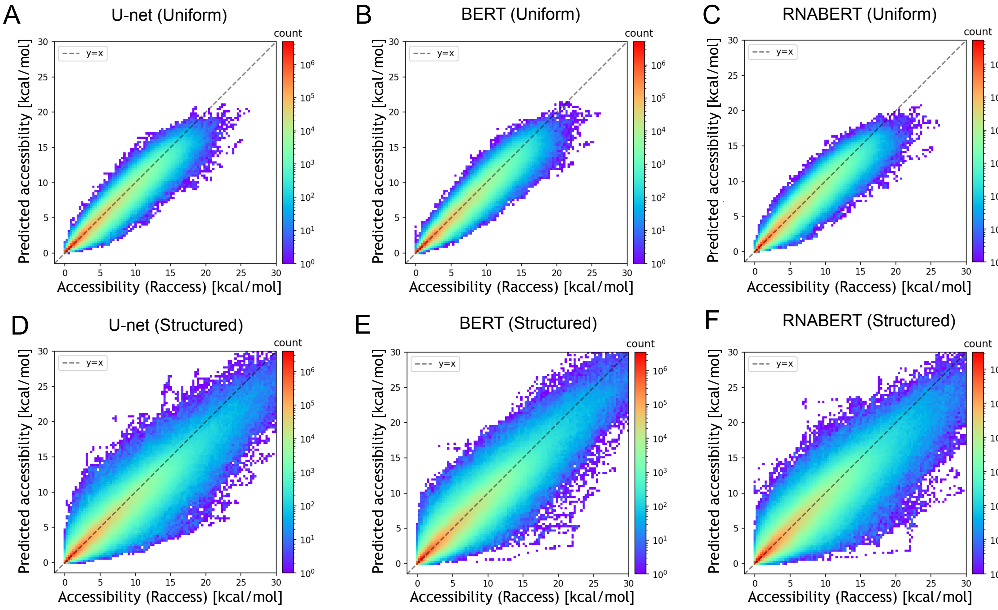


Fig. S1 Prediction performances for the simulation datasets. Accuracy of the uniform RNA dataset for (A) the U-net model, (B) the BERT model, and (C) the RNABERT model. Accuracy of the structured RNA dataset for (D) the U-net model, (E) the BERT model, and (F) the RNABERT model. The x and y axes represent the accessibility calculated by Raccess and predicted accessibility, respectively. The color bar representing the counts is displayed using a log scale.

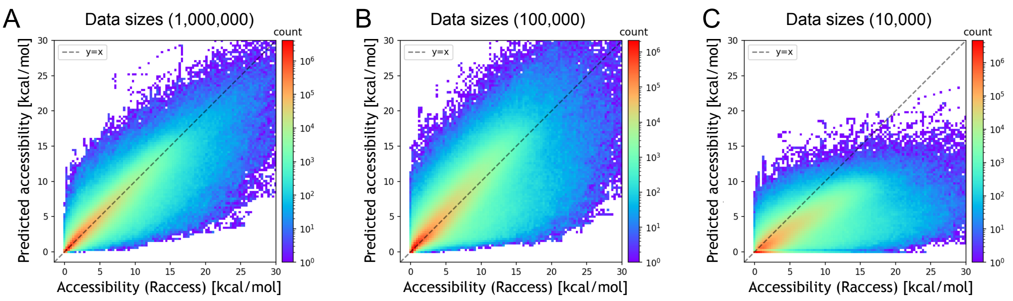


Fig. S2 Influence of the training data sizes on the prediction performances. The data sizes were (A) 1,000,000, (B)100,000, and (C) 10,000. The x and y axes represent the accessibility calculated by Raccess and predicted accessibility, respectively. The color bar representing the counts is displayed using a log scale.

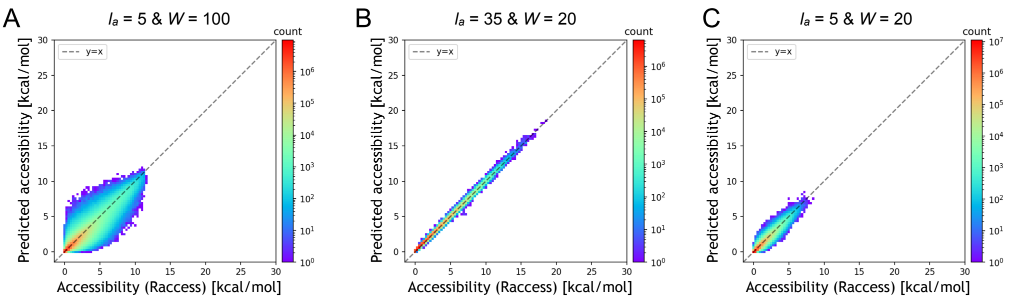


Fig. S3 Influence of the parameters $l_a$ and $W$ on the prediction performances. (A) $l_a = 5$ and $W = 100$. (B) $l_a = 35$ and $W = 20$. (C) $l_a = 5$ and $W = 20$. The x and y axes represent the accessibility calculated by Raccess and predicted accessibility, respectively. The color bar representing the counts is displayed using a log scale.

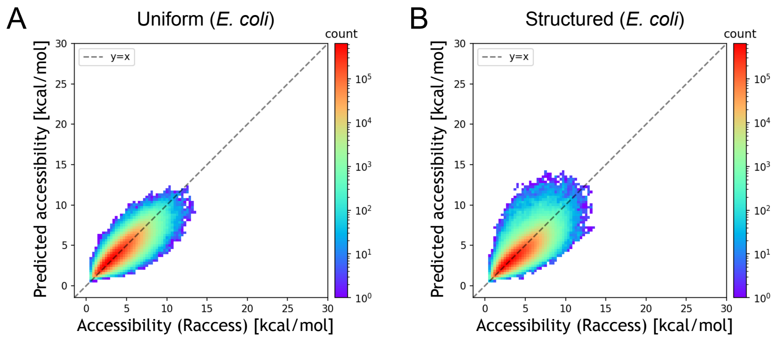


Fig. S4 Prediction accuracy for the *E.coli* synthetic RNA datasets of the predictive model trained on (A) the uniform RNA dataset and (B) the structured RNA dataset. The x and y axes represent the accessibility calculated by Raccess and predicted accessibility, respectively. The color bar representing the counts is displayed using a log scale.

### Supplementary Tables

Table S1 Architecture of the FCN model

| module | parameter | shape | notes |
|---|---|---|---|
| input | | (440) | |
| embedding | dim = 120 | (120, 440) | |
| 1D CNN BN Mish | chnnel=120, kernel=9, padding=4 | (120, 440) | |
| 1d CNN BN ReLU | chnnel=120, kernel=5, dilation=3, padding=6 | (120, 440) | × 40 |
| 1D CNN BN Mish | chnnel=120, kernel=9, padding=4 | (120, 440) | |
| 1d CNN BN ReLU | chnnel=1, kernel=9, padding=4 | (1, 440) | |
| output | | (440) | |

Table S2 Architecture of the U-net model

| module | parameter | shape | notes |
|---|---|---|---|
| nput | | (440) | |
| embedding | dim = 120 | (120, 440) | |
| 1d CNN BN ReLU | chnnel=240, kernel=5, padding=2, stride = 2 | (240, 220) | save as X |
| 1d CNN BN ReLU | chnnel=360, kernel=5, padding=2, stride = 2 | (360, 110) | save as Y |
| 1d CNN BN ReLU | chnnel=480, kernel=5, padding=2, stride = 2 | (480, 55) | save as Z |
| 1d CNN BN ReLU | chnnel=480, kernel=5, dilation=3, padding=6 | (480, 55) | save as Z |
| 1d CNN BN ReLU | chnnel=1, kernel=9, padding=4 | (480, 55) | × 35 |
| 1d transposedCNN BN ReLU | chnnel=360, kernel=5, padding=2, stride = 2 | (360, 110) | +Z |
| 1d transposedCNN BN ReLU | chnnel=240, kernel=5, padding=2, stride = 2 | (240, 220) | +Y |
| 1d transposedCNN BN ReLU | chnnel=120, kernel=5, padding=2, stride = 2 | (120, 440) | +X |
| 1D CNN BN Mish | chnnel=120, kernel=9, padding=4 | (120, 440) | |
| 1d CNN BN ReLU | chnnel=1, kernel=9, padding=4 | (1, 440) | |
| output | | (440) | |

Table S3 Architecture of the BERT model

| module | parameter | shape | notes |
|---|---|---|---|
| input | | (440) | |
| BERT | attention heads=12, hidden layers=6 | (120, 440) | |
| 1D CNN BN Mish | chnnel=120, kernel=9, padding=4 | (120, 440) | |
| 1d CNN BN ReLU | chnnel=1, kernel=9, padding=4 | (1, 440) | |
| output | | (440) | |

Table S4 Architecture of the RNABERT model

| module | parameter | shape | notes |
|---|---|---|---|
| input | | (440) | |
| BERT | attention heads=12, hidden layers=6 | (120, 440) | pretrained model |
| 1D CNN BN Mish | chnnel=120, kernel=9, padding=4 | (120, 440) | |
| 1d CNN BN ReLU | chnnel=1, kernel=9, padding=4 | (1, 440) | |
| output | | (440) | |

Table S5 Influence of the training data sizes on the prediction performances

| Data sizes | NMSE | Spearmans' $\rho$ |
|---|---|---|
| 10,000 | 1.6998 | 0.8601 |
| 100,000 | 0.6762 | 0.9449 |
| 1,000,000 | 0.3892 | 0.9688 |
| 10,000,000 | 0.1148 | 0.9876 |

Table S6 Influence of the parameters $l_a$ and $W$ on the prediction performances

| Parameters | NMSE | Spearmans' $\rho$ |
|---|---|---|
| $l_a = 5, W = 20$ | 0.0406 | 0.9744 |
| $l_a = 5, W = 100$ | 0.1396 | 0.9706 |
| $l_a = 35, W = 20$ | 0.0050 | 0.9995 |
| $l_a = 35, W = 100$ | 0.1148 | 0.9876 |
